# Exploring the Diversity of *Bacillus* whole genome sequencing projects using Peasant, the Prokaryotic Assembly and Annotation Tool

**DOI:** 10.1101/132084

**Authors:** Jonathon Brenner, Laurynas Kalesinskas, Catherine Putonti

**Affiliations:** Department of Computer Science, Loyola University Chicago, 820 N Michigan Avenue, Chicago, IL USA 60611; Department of Biology, Loyola University Chicago, 1032 W Sheridan Road, Chicago, IL USA 60660; Bioinformatics Program, Loyola University Chicago, 1032 W Sheridan Road, Chicago, IL USA 60660; Department of Microbiology and Immunology, Stritch School of Medicine, Loyola University Chicago, Maywood, IL, USA 60153

**Author notes:** Author Email Addresses: JB, LK, CP.

**Keywords:** genome assembly, genome annotation, automated pipeline, Bacillus comparative genomics

## Abstract

**Background:** The persistent decrease in cost and difficulty of whole genome sequencing of microbial organisms has led to a dramatic increase in the number of species and strains characterized from a wide variety of environments. Microbial genome sequencing can now be conducted by small laboratories and as part of undergraduate curriculum. While sequencing is routine in microbiology, assembly, annotation and downstream analyses still require computational resources and expertise, often necessitating familiarity with programming languages. To address this problem, we have created a light-weight, user-friendly tool for the assembly and annotation of microbial sequencing projects.

**Results:** The Prokaryotic Assembly and Annotation Tool, Peasant, automates the processes of read quality control, genome assembly, and annotation for microbial sequencing projects. High-quality assemblies and annotations can be generated by Peasant without the need of programming expertise or high-performance computing resources. Furthermore, statistics are calculated so that users can evaluate their sequencing project. To illustrate the computational speed and accuracy of Peasant, the SRA records of 322 Illumina platform whole genome sequencing assays for *Bacillus* species were retrieved from NCBI, assembled and annotated on a single desktop computer. From the assemblies and annotations produced, a comprehensive analysis of the diversity of over 200 high-quality samples was conducted, looking at both the 16S rRNA phylogenetic marker as well as the *Bacillus* core genome.

**Conclusions:** Peasant provides an intuitive solution for high-quality whole genome sequence assembly and annotation for users with limited programing experience and/or computational resources. The analysis of the *Bacillus* whole genome sequencing projects exemplifies the utility of this tool. Furthermore, the study conducted here provides insight into the diversity of the species, the largest such comparison conducted to date.

## BACKGROUND

Modern sequencing technologies have facilitated comprehensive characterization of genomes from a vast array of species, significantly expanding our view of genetic diversity on Earth [1]. With the advent of second- and third-generation sequencing platforms, the capabilities of DNA sequencers far surpassed that of the decades prior. The ever-decreasing cost of sequencing has spurred the transition of genome sequencing projects from a select few facilities to individual laboratories, and even curricular activities (e.g. [2,3]). With this transition in technology and increase in throughput, however, a major overhaul in the bioinformatic solutions for genome assembly was required.

Software tools for the processes of genome assembly and annotation are numerous. In addition to reference guided assembly strategies, *de novo* assemblers have been created including, e.g. Abyss [4], Velvet [5], SOAPdenovo [6], and SPAdes [7]. Additionally, newer long read platforms, such as PacBio and Nanopore, have incited the development of a new class of genome sequence assemblers, such as Canu [8] and miniasm [9]. In parallel, additional software solutions have been developed for read quality filtering (e.g. FASTQC [10], Trimmomatic [11], and Sickle [12]), assembly quality assessment (e.g. QUAST [13] and REAPR [14]), and genome scaffolding (e.g. SSPACE [15] and SCARPA [16]). In contrast to the previously mentioned assembly tools, which are largely local command line solutions, annotation solutions are more prevalent as web-based solutions, e.g. NCBI’s Prokaryotic Genome Annotation Pipeline [17], BASys [18], RAST [19], IGS [20], IMG [21], and Genix [22]. Nevertheless, the standalone solution Prokka [23] is increasingly popular; the ability to annotate genomes locally has the benefit of protecting sensitive data and often completes annotations more rapidly than web-based tools. Moreover, a local solution provides users greater control on how annotations are performed. While this is a far from an inclusive list of software solutions that have emerged, it is a testament to the fact that molecular biology is increasingly integrating computational approaches [24].

While DNA sequencing is currently more accessible, well-trained individuals that can conduct the bioinformatics workflow of a genome sequencing project are not as plentiful. Several pipelines have been developed to facilitate the process of assembly and quality control (QC), including A5 [25] and RAMPART [26]. Recently, tools have also been built to automate the entire processes of genome assembly and annotation, e.g. MEGAnnotator [27], MyPro [28], iMetAMOS [29], and the web-based pipeline PATRIC [30]. These all-in-one solutions integrate existing software (such as the assemblers, QC tools, and annotation tools previously listed) in a single tool. Thus, users only need knowledge concerning the single tool; the requirements and terminology of the individual software within are concealed from the user. While easy to use, installation of the individual components packaged in the pipeline can present significant challenges and/or require access to high performance computing resources.

Herein, we present a new automated pipeline for bacterial genome assembly and annotation called Peasant (available at https://github.com/jlbren/peasant). Raw reads, supplied by the user, are filtered for quality and assembled by Peasant; users can select from available filters to automate the quality control of their assemblies. This assembly is then annotated, identifying protein coding genes, tRNAs, and rRNAs using a local database. The motivation behind the development of this tool is three-fold. First, we saw a need for a robust, yet user-friendly solution for assembly, annotation, and reporting with automated QC options. As such, available assembly and annotation software were evaluated for their ease of use and precision. Peasant integrates several QC steps throughout the process. Second, while web-based solutions provide a resource for users with limited computational resources, they may limit the user’s ability to fine-tune the process for their study. Thus, Peasant was designed to include flexibility while necessitating only resources now commonplace in laboratories. Third, raw sequence data provides an unbiased representation of the organism sequenced: published assemblies may be produced by outdated tools or generated to meet criteria often unknown to the downstream user. Comparative genomic studies may thus benefit by returning to raw data. Peasant provides a means to feasibly process numerous genomes in a uniform manner. To illustrate the utility of Peasant, the SRA records for 322 *Bacillus* whole genome sequencing projects were retrieved from NCBI [31]. Each record was analyzed by Peasant, facilitating downstream analysis of the diversity of *Bacillus* species.

## IMPLEMENTATION

The Prokaryotic Assembly and Annotation Tool - Peasant - automates assembly and annotation using existing tools as well as novel functionality developed in Python. Figure 1 illustrates this process.

**Figure 1.**
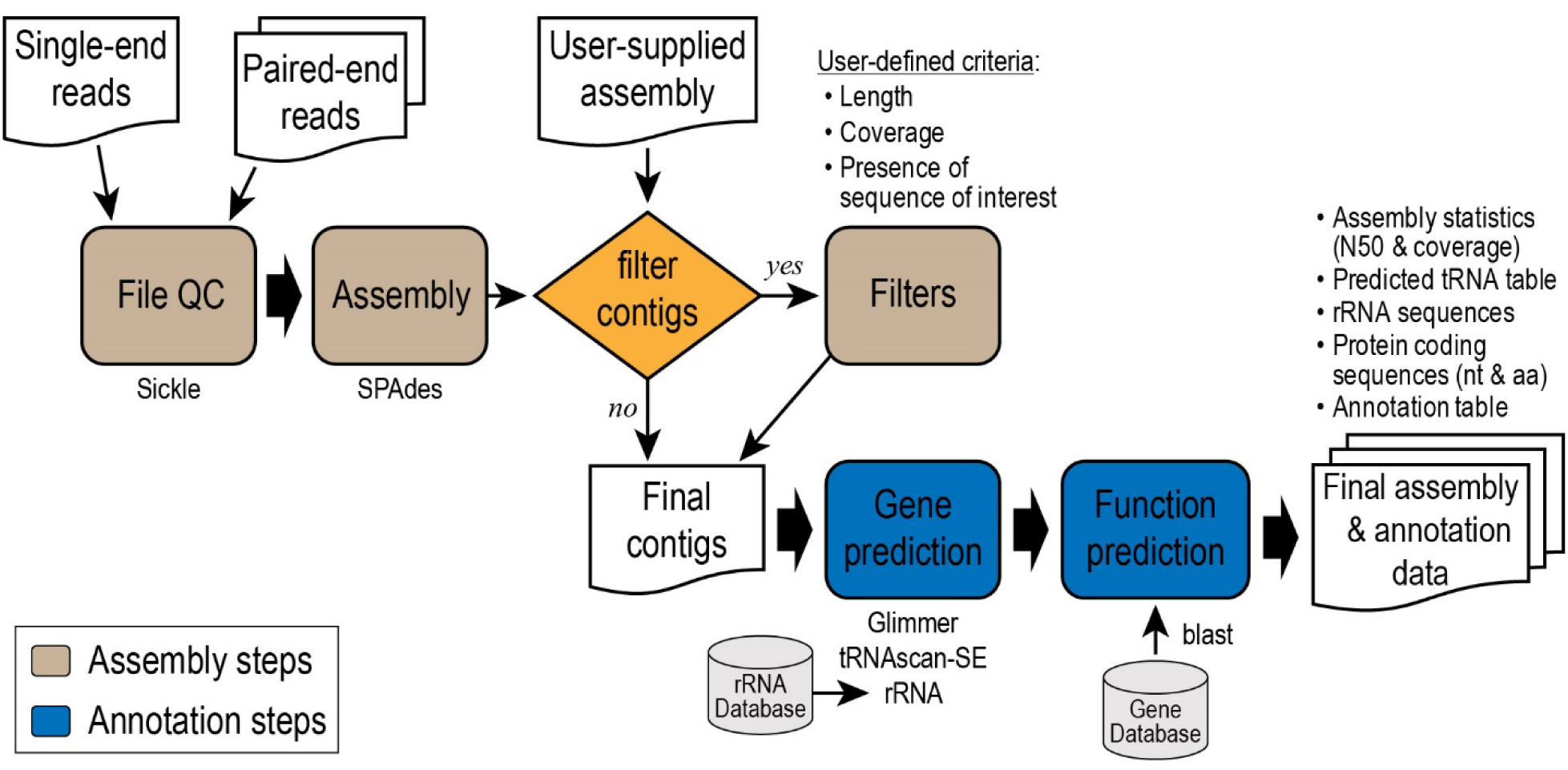
Schematic of Peasant process.

***Assembly Steps:*** Sequencing reads, either single or paired-end, are first processed and read QC is conducted using Sickle [12]. Assembly is next performed using SPAdes [7]. SPAdes was selected here as it is (1) frequently used in WGS studies, (2) it often outperforms other assemblers on microbial genomic projects (e.g. [27]), and (3) performs well with feasible demands on RAM. Moreover, current versions of SPAdes include the capability to conduct hybrid assemblies of reads from long and short read technologies. The code was developed to easily accommodate other assemblers, and future versions of this tool are anticipated to reflect new/additional tools in the field if they can provide quality assemblies at low cost (in terms of memory and time requirements). SPAdes is executed with the ‘--only-assembler’ option with word size k=33, 55, 77, 99 and 127. Alternatively, users can supply their own genome assembly file, removing the assembly process. Assemblies can then be filtered based upon user supplied criteria, including length and coverage. Table 1 lists these filter parameters.

**Table 1.** Filter parameters for Peasant.

| Parameter | Parameter Description | Example |
| --- | --- | --- |
| -m, --min_contig_size | Minimum contig size | -m 1000 |
| -M, --max_contig_size | Maximum contig size | -M 100000 |
| -c, --min_coverage | Minimum coverage value<br>(calculated by BBMAP [32]) | -c 100 |
| -cov, --min_SPAdes_cov | Minimum SPAdes cov value<br>(SPAdes assemblies only) | -cov 10 |

***Annotation Steps:*** Contigs are next examined, identifying rRNA, tRNA, and protein coding regions. 5S, 16S, and 23S rRNA regions are detected through BLASTn queries to 5S, 16S, and 23S blast sequence databases. The three rRNA databases, included with the download of Peasant (available through the github repository), were curated from the RNAmmer Server [33]. tRNA sequences are predicted using tRNAscan-SE [34]. Coding regions are predicted by running Glimmer with the g3-iterated script [35]; this script creates a training set from the genome assembly, builds an Interpolated Context Model (ICM) from the training sequences, runs Glimmer, creates a position weight matrix from the predicted sequences, and runs Glimmer again to generate the final coding region predictions. These predicted protein coding sequences are subsequently assigned functionality by BLAST queries to an annotated gene database. Users can specify the threshold for ascertaining homologous genes by specifying the minimum percent identity, query coverage, and/or blast bitscore; otherwise, default values are used: 70%, 70%, and 50, respectively. A precomputed gene database, created from all archived RefSeq bacterial genomes [36,37], is available from the github repository. Users can also create their own custom database for annotations by using the script make_Peasant_db.py (also available through the Peasant github repository). This script will produce Peasant formatted databases from user-supplied sequence (ffn format) and annotation information (ptt format).

***Outputs:*** In addition to the assembled genome file, several other files are generated by the tool, as indicated in Figure 1. The log file includes assembly statistics generated by Peasant, as well as the parameters used in the analysis, sequence file names, and commands implemented when calling external programs. Identified rRNA sequences, tRNA predictions, and protein coding sequences (as both nucleotide and amino acid sequences) for the assembled genome are also written to file. Finally, a CSV file is generated listing annotation details, including, the protein coding region’s location within the contig sequence, the predicted gene name, and product information.

***Tool specifics:*** This tool was developed in Python 2.7 and can be run on a UNIX/Mac OSX system through the terminal. As existing software tools are integrated into the tool, it is required that these tools are installed and included in the system’s PATH environmental variable. Inclusion of all necessary packages is automatically checked by Peasant upon execution and the user will be notified of any missing components. The tool’s dependencies include the BioPython Package [38], Sickle [12], SPAdes [7], Glimmer [35], BLAST+ [39] and BBMAP [32]. The documentation for Peasant includes links to these tools and their respective installation instructions and is available with the aforementioned scripts at https://github.com/jlbren/peasant. In addition to parameters required for executing the assembly and annotation of the sequencing reads, Peasant includes a parameter in which the user can specify the number of threads to use during BLAST searches. This parameter is integrated into calls to both the SPAdes assembler and annotation process and thus can significantly expedite the execution of Peasant.

## RESULTS AND DISCUSSION

### The Peasant software

Peasant is specifically tailored for whole genome assembly and annotation of an isolated bacterium. Execution of Peasant is through the command line in which users indicate their read files, database for annotation, and output location. This automated process provides feedback to the user via the log file which can indicate a poor-quality sequencing run and/or DNA prep. In addition, users can specify parameter values to filter their assembly (Table 1). These filters can remove low coverage contigs which may represent sequences of contaminants. Similarly, the existing pipelines MEGAnnotator [27] and PATRIC [30] include assembly filters for size and coverage; neither MyPro [28] nor iMetAMOS [29] include such functionality. The simplicity of Peasant provides mechanisms for even non-technical users to analyze their data: users can easily test different filters on assembled contigs and evaluate the quality of their sequencing data. Once an assembly is generated, users do not need to run the tool from start to finish to test filters or rerun annotations; Peasant can also accept a FASTA or multi-FASTA format assembly as a user input and thus bypass the assembly step of the pipeline. Peasant’s run-time is dependent upon the number of input contigs and the size of the annotation database. Using the multi-threading parameter, this can be expedited (dependent upon the number of threads supported by the user’s machine).

### Case Study: Assembly and annotation of publicly available *Bacillus* spp. whole genome sequencing projects

All *Bacillus* SRA files were identified using NCBI’s SRA Run Selector [40]. Projects self-identified as ‘WGS’ generated using an Illumina instrument were selected and downloaded. Metadata for each sample was manually inspected to verify it was for a *Bacillus* species and from single isolates. Supplemental Table 1 lists the 322 projects retrieved. Peasant then processed each sample with a single filter specified, which removed contigs less than 1000 bp in length. As part of the Sickle read QC process, we restricted trimmed reads to be greater than or equal to 100 nucleotides in length. This read length threshold automatically removed 14 samples (as these studies generated reads <100) as well as an additional 59 samples that did not meet this threshold post-trimming. Read QC is a critical step in genome assembly; only the existing pipeline MEGAnnotator [27] includes read QC as part of their automated process. Thus, in total, 249 *Bacillus* isolates were fully processed. Figure 2 provides an overview of the data sets, which included both single and paired-end reads through QC and assembly. Full statistics for each individual sample, including the number of predicted protein coding genes, rRNAs, and tRNAs can be found in Supplemental Table 2.

**Figure 2.**
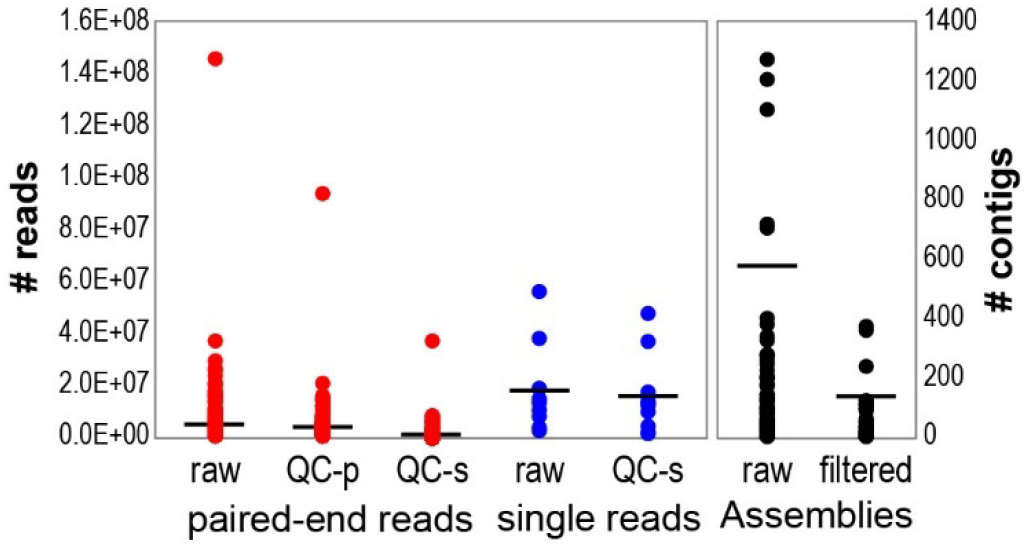
Overview of the 249 *Bacillus* samples processed through read QC and assembly. Each dot represents a single sample and the median number of reads or contigs across all samples is shown by a bar.

The uniform processing of the *Bacillus* samples facilitates subsequent comparative analyses. For each sample processed, Peasant generates a file containing the rRNA (5S, 16S and 23S) gene sequences identified. The 16S rRNA gene sequences were extracted from each sample; sequences with a length less than 1000 nucleotides were omitted from further analysis. It is worth noting that some samples did not contain a recognized 16S rRNA sequence and/or a partial sequence and thus were not included in subsequent analyses. In total 218 16S rRNA gene sequences were aligned with 30 *Bacillus* RefSeq [36] 16S rRNA gene sequences (retrieved from NCBI, listed in Supplemental Table 3) and two out-group sequences – *Paenibacillus polymyxa* SC2 (NC_014622) and *Lactobacillus casei* BLS23 (NC_010999). This alignment was used to derive a phylogenetic tree using the tool FastTree [41]. *L. casei* was specified as the root of the tree shown in Figure 3. Branches without a label are representative of individual samples assembled in this study.

**Figure 3.**
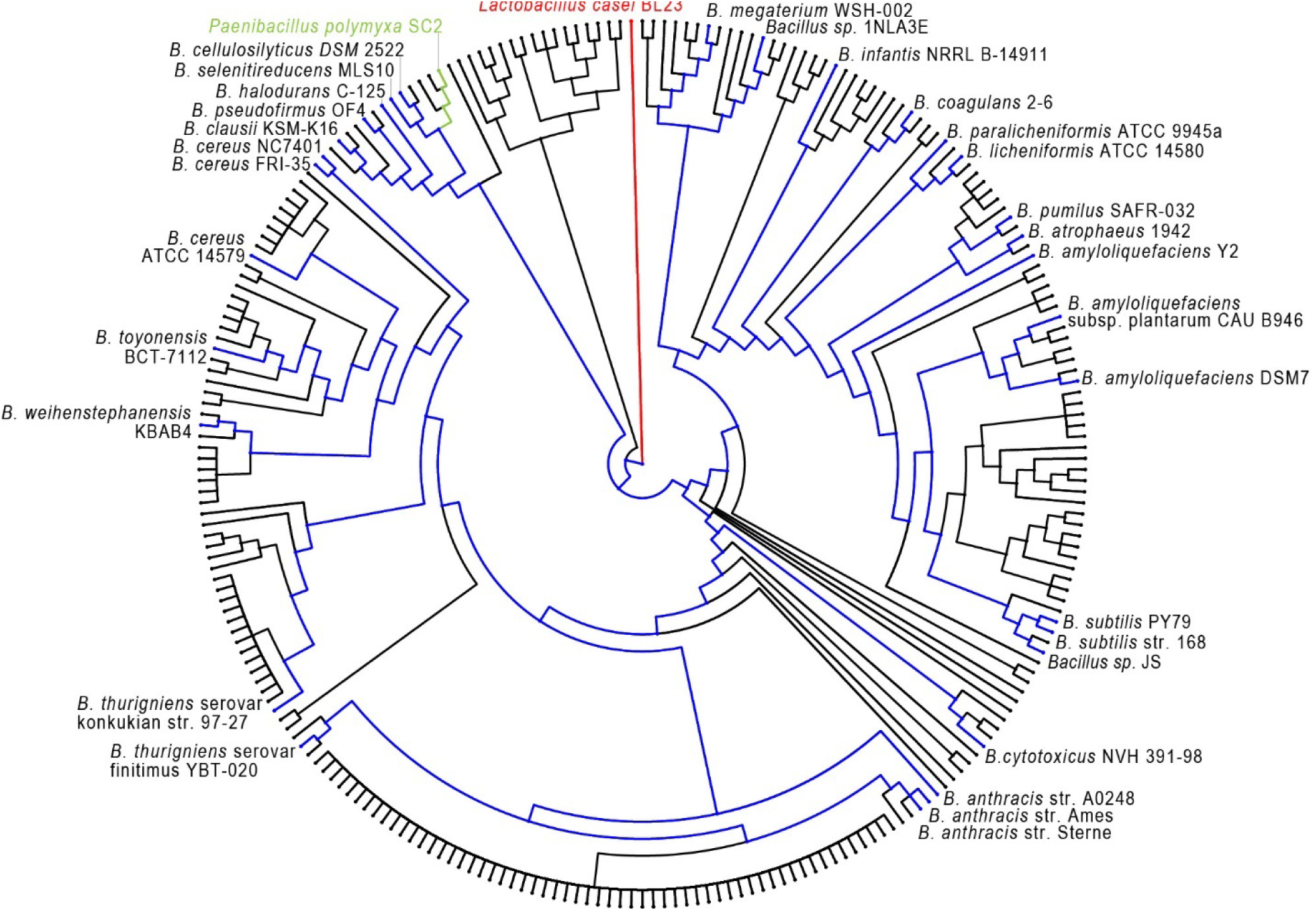
16S rRNA gene sequence phylogeny of *Bacillus* sequencing projects and *Bacillus* spp. RefSeq representatives (indicated in blue). The 16S rRNA gene sequences for *L. casei* (red) and *P. polymyxa* (green) are included as outgroups. The tree was derived using FastTree [41], an approximate maximum-likelihood method, with support values (not shown), and visualized using Phylowidget [42].

As previous studies have found, distinguishing bacilli species via molecular methods (such as 16S rRNA) is not possible for some taxa, e.g. the *B. cereus* sensu lato group which includes *B. cereus*, *B. thuringiensis*, *B. anthracis*, *B. mycoides*, *B. pseudomycoides*, and *B. weihenstephanensis* [43-46]. These individual species were not found to be monophyletic in the 16S rRNA tree of Figure 3. Of interest is the clade shown in Figure 3 between the two outgroup species labels. This branch is separate from both the outlier *L. casei* and the rest of the bacilli sequences. The 16 sequences within this clade were selected and queried via NCBI’s BLAST web interface against the ‘16S ribosomal RNA sequences (Bacteria and Archaea)’ database. All 16 sequences produced hits to taxa other than *Bacillus* (Supplemental Table 4), including hits to 16S rRNA gene sequences from *Clostridium*, *Francisella*, and *Methylobacterium* species. Five of these 16 sequences are from samples in which there is only one recognized 16S rRNA sequence; thus, based on 16S rRNA sequence alone, we conclude that the isolate sampled is not a *Bacillus* species. The Peasant annotation from the remaining 11 sequences within this clade are from samples in which more than one 16S rRNA gene sequence was identified. This 16S sequence was *Bacillus* in origin (details listed in Supplemental Table 4) suggesting the presence of a contaminant within the samples sequenced. This is certainly the case for SRA record SRR2120204 with one 16S sequence producing a hit to

*Citrobacter murliniae* strain CDC 2970-59, and with a second 16S sequence in the same clade as *B. cereus* ATCC 14579 in Figure 3. The assembly produced for the SRR2120204 read set exceeds 13Mbp, which well exceeds the genome size of either taxa.

While there are presently >1500 assemblies in GenBank for *Bacillus* species (https://www.ncbi.nlm.nih.gov/assembly/?term=txid1386[Organism:exp]), the number of whole genome sequencing projects for *Bacillus* species exceeds 2000. The ability to process raw sequencing data quickly, easily, and uniformly from many studies, such as the *Bacillus* spp. examined here, provides an invaluable tool for researchers conducting comparative genomics studies. Given the wealth of genomic sequences available we can begin to explore the evolutionary history of the *Bacillus* genus. While this can be approximated via analyses of, e.g., the 16S rRNA gene (Figure 4), identification and phylogenetic analysis of the ‘core genome’ provides a significantly more robust measure. Prior studies have found at least 600 genes within the core genome of the *B. cereus* sensu lato group [47,48]. From the examination of 20 *Bacillus* genomes, including taxa outside of the *B. cereus* sensu lato group, 814 orthologous genes were identified as the core genome of the genus [49]. Looking here at a larger collection of sequences than previously considered, we identified the *Bacillus* core genome from the predicted coding regions generated by Peasant. (Core genome analysis considered 231 high-quality assemblies; samples were identified as ‘high-quality’ assemblies based upon their 16S rRNA sequences, number of predicted genes, and genome size (Supplemental Table 1).) Thirty genes were identified within all assembled isolates and 310 genes were identified within over 95% of the isolates (Supplemental Table 5). As shown in Figure 4, there is a large core genome (5023 genes) found within ∼70% of the genomes examined; this reflects the over-representation of taxa within the dataset belonging to the *B. cereus* sensu lato group. Inclusion of more distant relatives to this group reduces the number of genes associated with the genus’ core genome.

**Figure 4.**
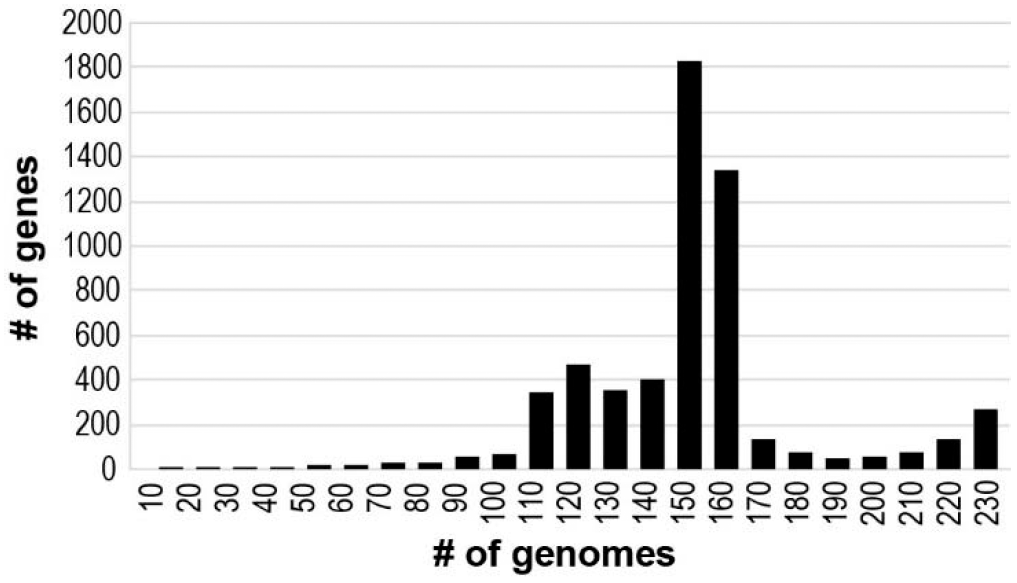
Evaluation of the core genome within 231 *Bacillus* isolates.

## CONCLUSIONS

Peasant provides an expedient way to take raw sequencing reads and produce annotated assemblies. Its lightweight construction makes it feasible to conduct whole genome studies without high performance computing resources or programming expertise. The modular design of the tool permits the addition of new tools easily and future development of Peasant will expand options of assembly, QC, and annotation. With the consistent rise of whole genome sequencing efforts, the ability to expediently perform uniform processing provides an unbiased platform for bioinformatic analysis. The case study presented here, looking at over 200 individual *Bacillus* genomes, exemplifies the use of Peasant for large scale analyses. Our uniform processing of samples has identified samples with concerns of contamination and/or mixed communities as well as putative mislabeling. Furthermore, the examination of these *Bacillus* genomes is the largest study to date into the core genome of this genus.

## ACKNOWLEDGEMENTS

The authors would like to thank Dr. Alan Wolfe and Ms. Krystal Thomas-White for their feedback during development of this tool.

## Supplemental Table Legends

**Supplemental Table 1.** List of the 322 *Bacillus* sequencing SRA projects retrieved.

**Supplemental Table 2.** Assembly and annotation statistics for processed *Bacillus* sequencing SRA projects.

**Supplemental Table 3.** NCBI RefSeq 16S rRNA gene sequences included in phylogenetic analysis of assembled *Bacillus* isolates.

**Supplemental Table 4.** Details regarding samples from which 16S rRNA sequences from taxa other than *Bacillus* were identified.

**Supplemental Table 5.** Genes identified within all (Sheet 1) 231 high-quality *Bacillus* genome assemblies and within 95% (Sheet 2) of the high-quality *Bacillus* genome assemblies.

